# The chemokine receptor CXCR2 contributes to murine adipocyte development

**DOI:** 10.1101/339721

**Authors:** Douglas P Dyer, Joan Boix Nebot, Christopher Kelly, Laura Medina-Ruiz, Fabian Schuette, Gerard J Graham

## Abstract

Chemokines are members of a large family of chemotactic cytokines that signal through their receptors to mediate leukocyte recruitment during inflammation and homeostasis. The chemokine receptor CXCR2 has largely been associated with neutrophil recruitment. However, there is emerging evidence of roles for chemokines and their receptors in processes other than leukocyte migration. We have previously demonstrated that CXCR2 KO mice have thinner skin compared to wild type mice. Herein we demonstrate that this is due to a thinner subcutaneous adipose layer, as a result of fewer and smaller individual adipocytes. We observe a similar phenotype in other fat depots and present data that suggests this may be due to reduced expression of adipogenesis related genes associated with adipocyte specific CXCR2 signalling. Interestingly, this phenotype is evident in female, but not male, CXCR2 KO mice. These findings expand our understanding of non-leukocyte related chemokine receptor functions and help to explain some previously observed adipose-related phenotypes in CXCR2 KO mice.

## Introduction

Chemokines belong to a family of approximately 50 structurally related proteins that signal through their cognate G-protein coupled receptors (GPCRs) to mediate cellular migration and recruitment^1^. Chemokine receptors are expressed on leukocytes at rest and during inflammation and play an important role in the multi-step process of leukocyte migration from the circulation, across the endothelium and within tissues^2^. Chemokine activity is also fine-tuned by a subfamily of receptors called atypical chemokine receptors (ACKRs) which are stromally expressed and which regulate local chemokine presentation and availability^3^. Given their central role in leukocyte recruitment, chemokines and their receptors are integral to the immune response and thus inflammatory based diseases^4^. The chemokine system has therefore been a target for therapeutic intervention in inflammatory disease. However there has been limited success in this regard^5,6^ and this is thought to be at least partially due to our lack of basic understanding of the breadth and complexity of chemokine function^6^. Chemokines and their receptors also play a key role in homeostatic processes. For example the CXCR4 receptor and its ligand CXCL12 are critical to retention and maturation of haematopoietic stem cells in the bone marrow^7^ and other homeostatic chemokine receptors are central contributors to cellular address codes ensuring precise temperospatial leukocyte migration under resting conditions^6^. In addition to their involvement in regulating cellular migration in immune and inflammatory responses chemokines and their receptors, in association with the ACKRs, display evidence of significant pleiotropy contributing to the regulation of processes such as angiogenesis^8^, cellular proliferation^9^ and apoptosis^10^. Chemokines, their receptors and the ACKRs also play key roles in development including governing stem cell migration within the embryo and contributing to the regulation of branching morphogenesis^11,12^.

CXCR2 is a chemokine receptor that has largely been associated with expression, and function, on neutrophils during inflammatory responses. Along with its ligands CXCL1-3 and CXCL5-8, CXCR2 controls neutrophil release from the bone marrow^13^ and enables their recruitment to inflamed and infected sites^14,15^. An additional, non-canonical CXCR2 ligand, MIF, has also been identified and is thought to play a role in recruitment of myelo monocytic cells during atherosclerosis^16,17^. Given the role of neutrophils in propagating inflammatory disease CXCR2 is seen as a viable therapeutic target in a range of pathologies^18^, for example chronic obstructive pulmonary disease (COPD)^19^ and chronic pancreatic inflammation^20^. Furthermore, CXCR2 blockade is also suggested to be of potential use in cancer therapy due to its ability to prevent recruitment of pro-tumorigenic myeloid derived suppressor cells to the tumour microenvironment^21,22^.

We have previously demonstrated that adult female CXCR2 KO mice have thinner skin than wild type mice^23^. Here we present data explaining the basis for this phenotype. Specifically we show that CXCR2 KO female mice have smaller adipocytes in several different fat depots, possibly because of aberrant expression of CXCR2-regulated adipogenesis related genes^24^. Our findings may, at least in part, explain CXCR2 KO mouse protection from obesity-induced insulin resistance^25^. Overall the results presented broaden our understanding of the role of the chemokine system in adipose tissue regulation and add to the developmental processes in which chemokines play a role.

## Materials and Methods

### Mice

CXCR2 KO^15^ (backcrossed for at least 12 generations onto the C57/Bl6 background) and WT mice were bred in a specific pathogen free environment and fed a normal lab diet (ad libitum) in the animal facility of the Beatson Institute for Cancer Research and were used at the indicated age, juvenile (6 weeks) and adult (8-12 weeks), in accordance with the animal care and welfare protocols approved by the animal welfare and ethical review board at the University of Glasgow. All experiments were performed under the auspices of a UK Home Office project licence.

### Histology

Skin and fat depots were harvested from mice, fixed in formalin, processed and embedded in paraffin wax. 5 μm thick tissue sections were cut from these paraffin blocks, baked (65°C for 30 minutes) onto SuperFrost slides (Thermo Scientific), dewaxed in xylene, re-hydrated and stained with haematoxylin and eosin counterstain. Finally, slides were de-hydrated and mounted using DPX (Leica).

Myeloperoxidase (neutrophil) staining was undertaken by the Diagnostic services unit, School of Veterinary Medicine at the University of Glasgow. Astra blue (Mast cell) and MAC2 (macrophage) staining was undertaken as described in detail previously^23^. The indicated measurements were then taken from blinded samples using the Zen software platform (Zeiss).

### Adipocyte differentiation, stimulation and analysis

The 3T3-L1 fibroblast cell line (a gift from Professor Gwyn Gould, University of Glasgow) was differentiated as described elsewhere^26^, briefly cells were cultured in Dulbecco’s modified Eagles medium with 10% foetal calf serum at 37°C in 5% CO_2_. To induce differentiation, cells were grown to confluence before addition of differentiation medium (DMEM containing 10% FBS (Gibco), 5 μM Troglitazone (Tocris), 1 μg/ml Insulin (Sigma-Aldrich), 0.5 mM IBMX (Sigma-Aldrich) and 0.25 μM Dexamethasone (Sigma-Aldrich)). After 3 days culture, the media was changed to DMEM containing 10% FBS, 5 μM Troglitazone and 1 μg/ml Insulin. Following a further 3 days of culture, media was changed to DMEM containing 10% FBS only and then changed every 2 days. Differentiation was also undertaken in the presence of the CXCR2 inhibitor SB 332235 (Tocris) to determine its effect on this process. Oil Red-O staining was used to confirm differentiation. Media was aspirated and cells washed with phosphate buffered saline before fixation in 10% formalin solution for 1 hour. Monolayers were incubated with 60% Isopropanol for 2 minutes before staining with Oil Red-O solution for 5 minutes and washing with water. Finally cells were counterstained with haematoxylin and washed before imaging using an EVOS FL auto microscope (Invitrogen). Cell size and lipid area were calculated from acquired images using FIJI software^27^. Quantification of Oil Red-O staining was undertaken as described previously^19^, following fixation cells were washed with water and stained with 0.5% Oil Red-O (w/v) in isopropanol for 15 minutes. Cells were washed with water to remove excess dye before Oil Red-O was eluted with isopropanol and quantified (OD540) relative to blank wells. Western blot analysis of cell cultures lysed in RIPA buffer (Thermofisher) was undertaken as described elsewehere^19^ using an anti PPARγ antibody (Cell Signaling Technologies) and an anti GAPDH antibody (Cell Signaling Technologies).

### PCR and qRT-PCR analysis

Samples from fat depots were harvested and briefly stored in RNAlater (Invitrogen) at −80°C. RNA was then extracted from these samples using Qiazol and physically disruption by shaking with ball bearings in a Tissue Lyser LT (Qiagen) at 50 oscillations per second for 10 minutes. The fluid phase was transferred to a clean tube and centrifuged at 4°C, 10,000 RCF for 15 minutes. The fluid phase was aspirated and transferred to a fresh tube prior to standard RNA extraction using the miRNeasy extraction kit (Qiagen). Purified RNA (500 ng) was converted to cDNA using the high capacity RNA to cDNA kit (ThermoFisher Scientific). cDNA was analysed for CXCR2 expression using the indicated primers (Table 1), Q5 High-Fidelity DNA polymerase and DNTP mix (NewEngland Biolabs) using a standard thermocycler programme. The resulting products were ‘run’ on a 2% agarose gel containing ethidium bromide and the gel analysed using an Alpha Imager (Alpha Innotech), following electrophoresis.

**Table 1.**
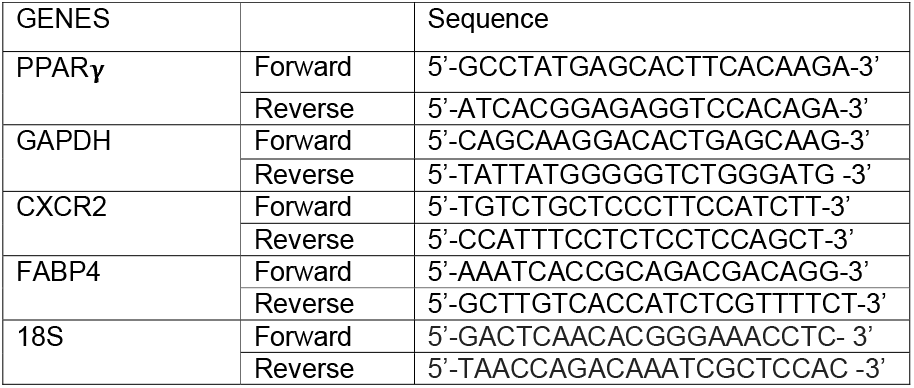
Primer sequences for qPCR analysis.

For qRT-PCR, cDNA was analysed using the indicated primers (Table 1) and SYBR Green FastMix (Quanta bio) in a QuantoStudio 7 flex machine (Life Technologies) and compared to standards for each gene, using the indicated primer sets^28^. Absolute copy number for each sample was then plotted relative to a house keeping gene control.

### Statistical analysis

Data sets containing only two groups were initially analysed for normal distribution (KS normality test) and equal variance (F test) using GraphPad Prism software. Normally distributed data with equal variance were analysed using an unpaired student’s t test, normally distributed data with unequal variance were analysed using an unpaired t test with Welch’s correction and non-normally distributed data with equal variance were analysed using a Mann Whitney test. Data sets containing more than two groups were analysed using a 1 way Anova with Tukey’s multiple comparison test. Where data were deemed significantly different (P<0.05) actual P values are provided in the figures from the indicated statistical test.

## Results

### Smaller adipocytes are associated with a thinner subcutaneous adipose layer in female CXCR2 KO, compared to wild type, mice

As reported in a previous study, and as shown in Figure 1, we observed that female CXCR2 KO mice have thinner skin^23^. Gross histological (Figure 1ai) and quantitative assessment (Figure 1aii) of sub-cutaneous adipose layer thickness in WT and CXCR2 KO female mice revealed that this reduced skin thickness is specifically associated with a thinner adipocyte layer and no significant differences in epidermal or dermal thickness were noted (data not shown). High power magnification revealed that the adipocytes in the CXCR2 KO mouse sub-cutaneous adipose layer were typically smaller than those in WT adipose tissues (Figure 1bi) and this was confirmed by measuring the size of individual adipocytes in the CXCR2 KO and WT mice which showed the CXCR2 KO adipocytes to be significantly (P=0.0021) smaller in size than the WT cells (Figure 1bii). In addition to reduction in size, the CXCR2 KO mouse adipose layer contained fewer cells as assessed by counting cells per field of view (Figure 1c). Overall therefore, these data show that female CXCR2 KO mice have a thinner adipose layer than WT mice and that this relates to smaller, and fewer, adipocytes within the CXCR2 KO adipose layer.

**Figure 1.**
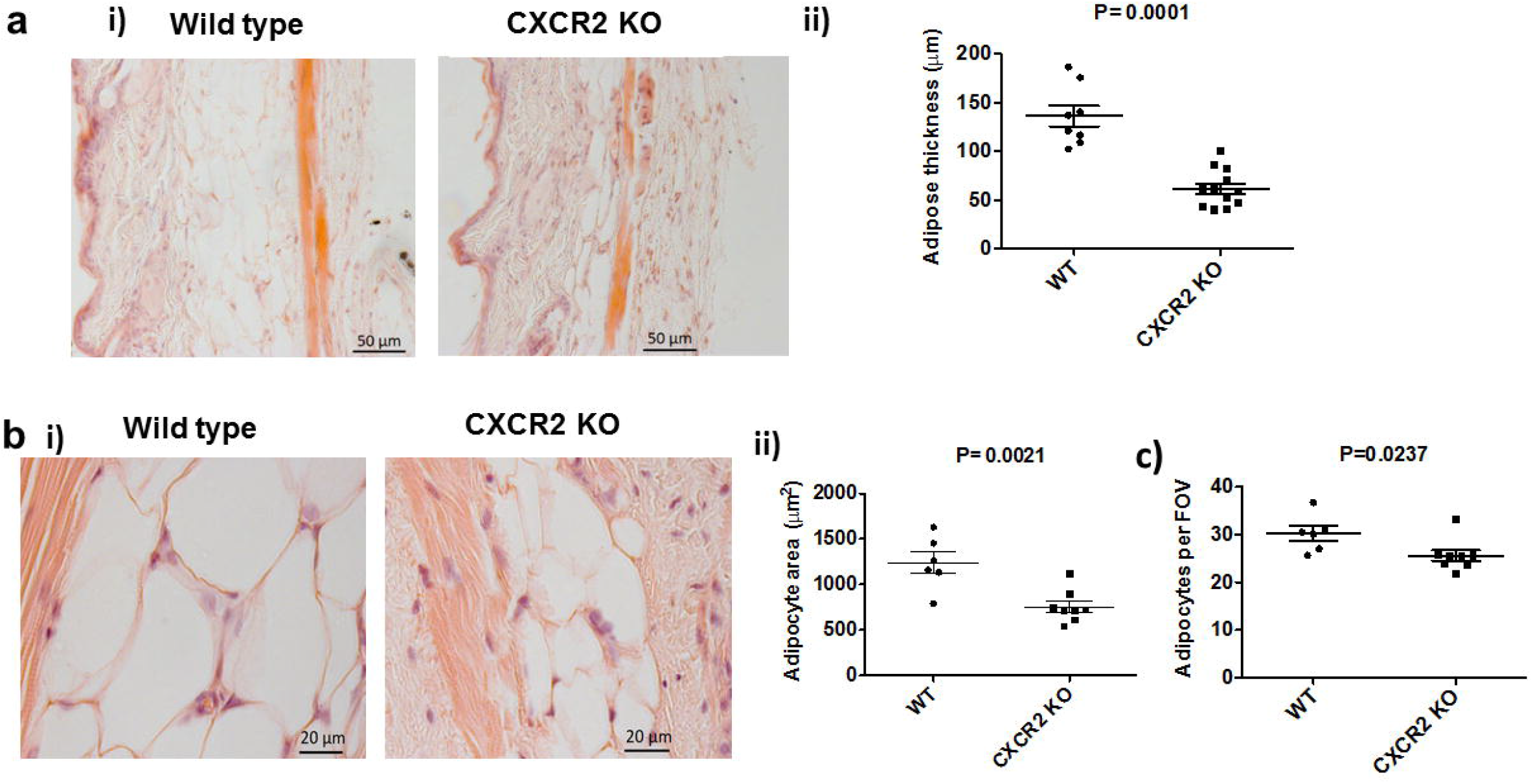
Female CXCR2 KO mice have a thinner subcutaneous adipose layer due to fewer and smaller adipocytes. Skin was dissected from adult female mice before processing and haematoxylin and eosin staining of sections from wild type and CXCR2 KO mice. (a) i) Brightfield microscopy was used to take images of the skin (scale bars-50μm) and ii) thickness of the adipocyte layer was systematically measured. (b) i) the subcutaneous adipose depot was further imaged at high magnification (scale bars-20μm) and ii) the size of individual adipocytes measured and expressed as μm^2^. (c) In addition, the number of individual adipocytes contained in the adipocyte layer (per field of view; FOV) was quantified. Data are plotted as mean (± SEM) from one experiment containing at least 5 mice in each group, representative of at least 2 separate experiments. Each symbol represents an individual mouse. Data were analysed with an unpaired t test with Welch’s correction (i), unpaired T test (ii) and a Mann Whitney test (iii).

### Adipocytes from multiple tissue sources are smaller in CXCR2 KO mice

Visual analysis of the anterior subcutaneous (tricep-associated), inguinal and perigonadal white adipose tissue reveals that reduced size of adipose depots in CXCR2 KO mice appears to be wide spread (Figure 2a). Closer histological analysis of tricep-associated, perigonadal and inguinal adipose depots in CXCR2 KO mice revealed, again, that smaller depots were associated with reduced adipocyte size (Figure 2b) and quantitative analysis revealed this to be a significant (P=0.0005, 0.0011 or 0.003, respectively) reduction (approximately 50%) at each of these tissue sites (Figures 2ci, 2cii and 2ciii). Interestingly, CXCR2 heterozygous null female mice display haploinsufficiency in adipocyte size in the tricep-associated and inguinal adipose depots, but not in perigonadal adipose tissue (Figure 2c).

**Figure 2.**
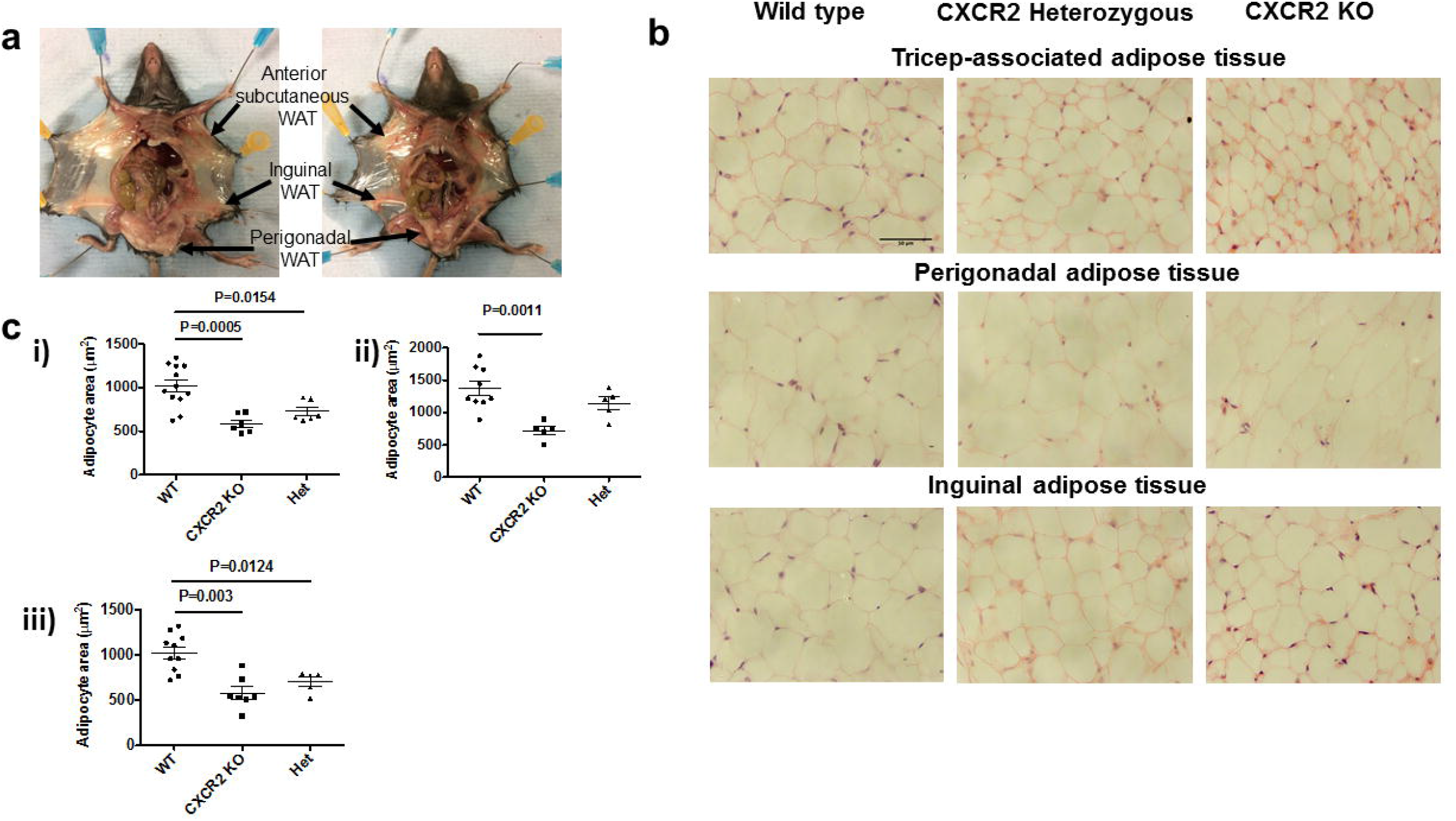
Adipocytes from multiple sources are smaller in CXCR2 KO female mice, compared to wild types. (a) Mice were dissected to allow visualization of fat depots at the indicated sites. (b) Adipose tissue was dissected, fixed and processed from each of these 3 areas and sections were stained with haematoxlyin and eosin (scale bars-50μm). (c) The area occupied by individual adipocytes was calculated and averaged for each individual mouse, data from each mouse was plotted for i) Tricep; ii) Perigonadal and iii) Inguinal sites, where each symbol represents an individual mouse. Data from two experiments are pooled and plotted as mean (± SEM). Data were analysed using a 1-way ANOVA with Tukey’s post hoc test.

### Reduced adipose tissue size is seen in female, but not male, CXCR2 KO mice

Our initial analyses were performed using female mice. To investigate any potential gender specificity of this phenotype, adult male CXCR2 KO mice were also analysed, in comparison to wild type mice. In contrast to female mice, CXCR2 KO male mice showed no significant differences in subcutaneous adipose thickness or adipocyte size in skin, inguinal or perigonadal adipose depots. This was apparent on gross histological assessment (Figure 3a) as well as on more detailed quantitative analysis which revealed no significant differences in sub-cutaneous adipose layer thickness (Figure 3b) and no significant difference in adipocyte size in inguinal or perigonadal sites (Figure 3ci and ii). Comparison with the data from analysis of adipose tissue and adipocytes in female mice (Figure 1) suggests that wild type females show generally increased subcutaneous adipose thickness and greater adipocyte size compared to males and that CXCR2 KO females have subcutaneous adipose tissue, and adipocytes, of comparable size to those in wild type males of the same age. Taken together these data suggest that CXCR2 plays a role in regulating adipocyte size in female, but not male, mice in resting adipose tissue.

**Figure 3.**
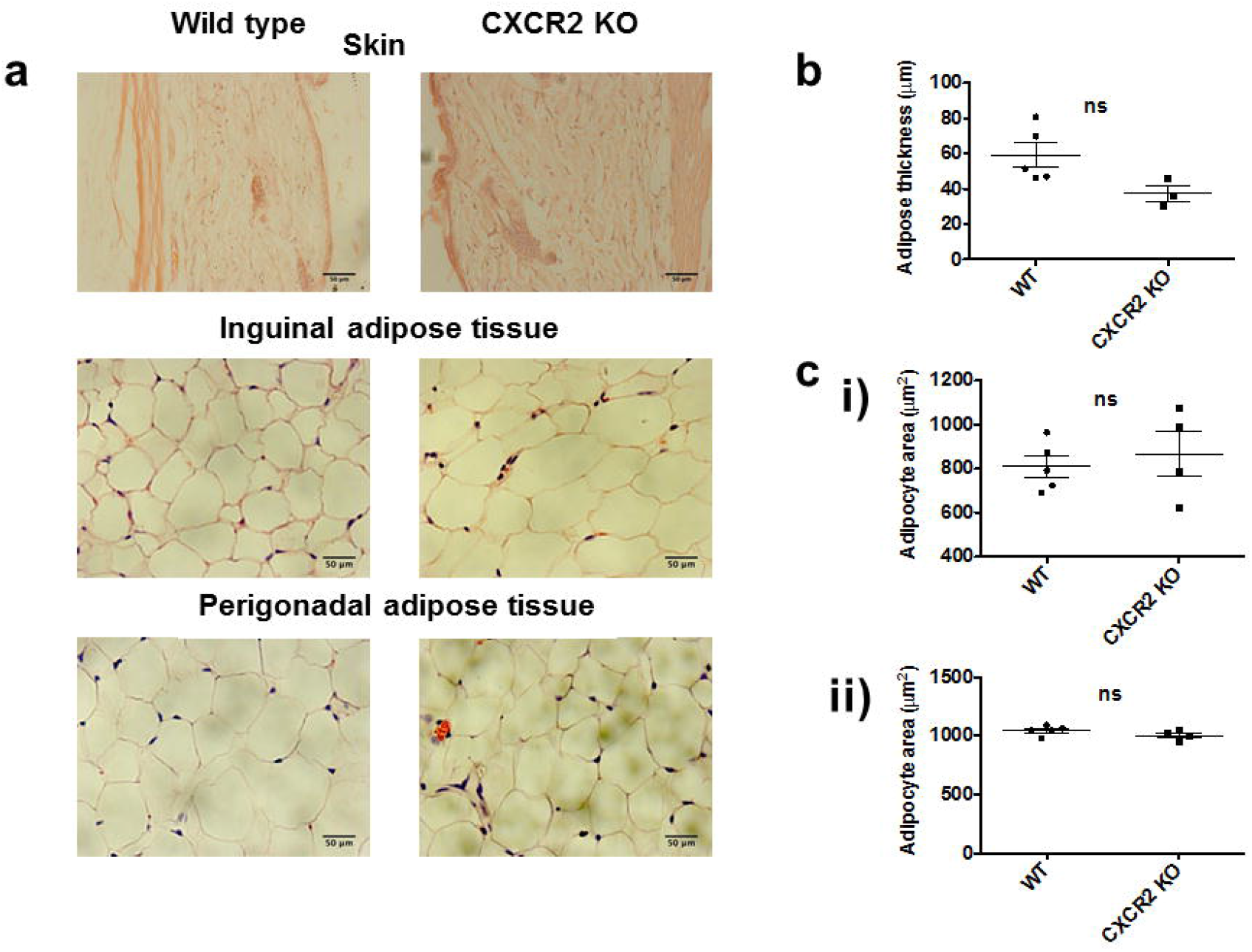
CXCR2 KO male mice have no significant change in adipocyte size compared to wild types. (a) Skin and adipose depots were dissected from adult male mice (>8 weeks) before processing and haematoxylin and eosin staining of sections from wild type and CXCR2 KO mice. Brightfield microscopy was used to take images of the skin, inguinal adipose or perigonadal adipose tissue (scale bars-50μm). (b) Adipose thickness and (c) individual adipocyte area for i) inguinal and ii) perigonadal sites were measured. Data are plotted as mean (± SEM), where each symbol represents data from an individual mouse and analysed using an unpaired t test, ns = not significant.

### The subcutaneous adipose layer from CXCR2 KO mice contains comparable levels of myelomonocytic cells to wild type mice

The cells traditionally associated with CXCR2 expression, and function, are neutrophils which are key players in the inflammatory response^18^. For this reason, and given previous studies describing a link between adipose tissue regulation, chemokine receptors (CXCR2, CCR2 or CX3CR1) and inflammation^29–31^, we determined the presence of neutrophils, mast cells and macrophages within the subcutaneous adipose layer of the skin of WT and CXCR2 KO female mice (Figure 4). As expected, myeloperoxidase staining clearly demonstrated the complete absence of neutrophils in the subcutaneous adipose tissue of CXCR2 KO mice (Figure 4a) and this was seen for both female and male (data not shown) mice. Furthermore Astra blue, and Mac 2, staining demonstrated no visual (Figure 4b and c) or quantitative (Figure 4d) changes in absolute numbers of mast cells and macrophages, respectively, between wild type and CXCR2 KO mice. In addition the ratio of mast cells, or macrophages, to adipocytes was unchanged between wild type and CXCR2 KO female mice (Figure 4e). Together, these data suggest that alterations in the levels of neutrophils, mast cells or macrophages do not explain the difference in subcutaneous adipocyte size between wild type and CXCR2 KO female mice.

**Figure 4.**
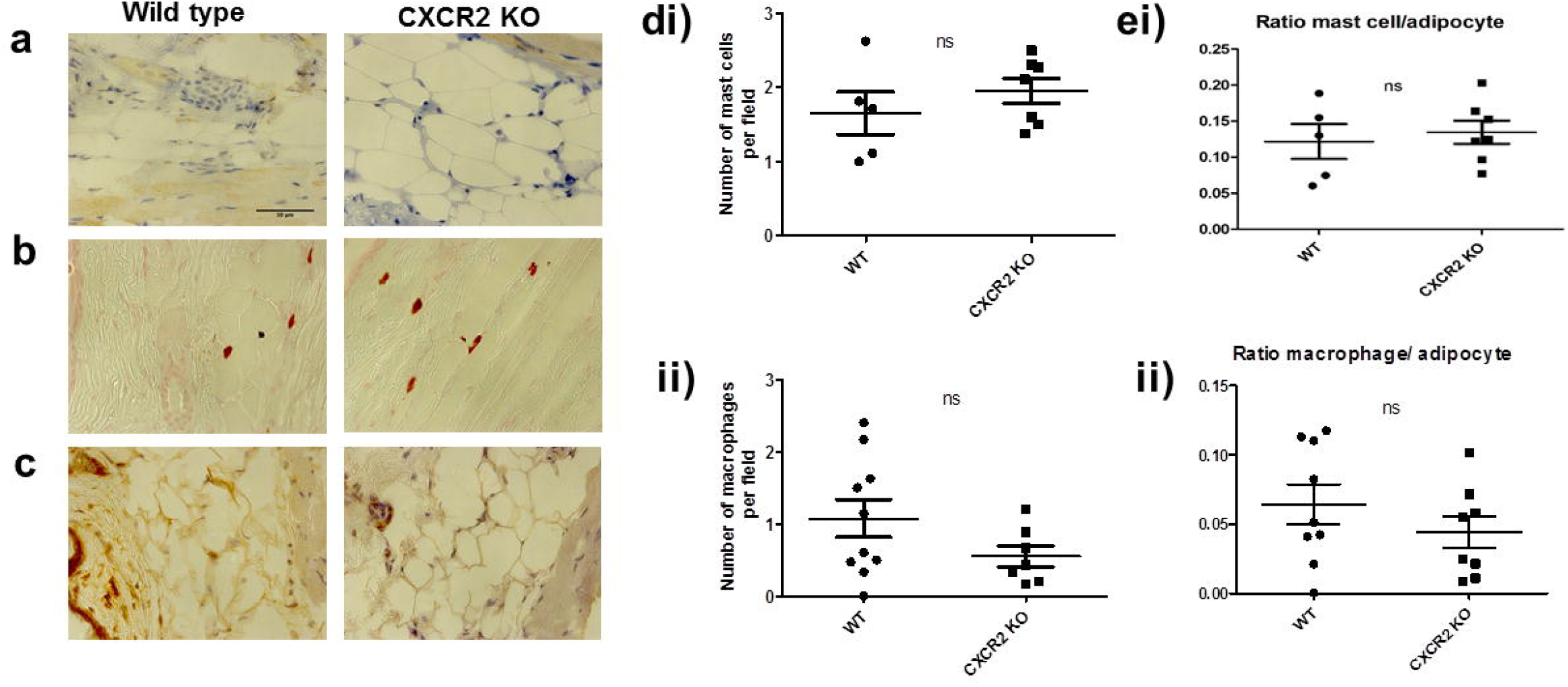
Resting CXCR2 KO mice show no differences in neutrophil, mast cell or macrophage numbers in subcutaneous fat. Skin was dissected from adult female mice before processing and staining (scale bars-50μm) to detect (a) neutrophils, (b) mast cells and (c) macrophages from wild type and CXCR2 KO mice. (d) The numbers of mast cells (i) and macrophages (ii) were quantified and plotted as the mean number per field of view. (e) i) Mast cell and ii) macrophage numbers were also expressed as ratio of cell number to adipocyte number. Data are plotted as mean (± SEM), where each symbol represents data from an individual mouse and analysed using an unpaired t test, representative of two separate experiments, ns = not significant.

### Adipocytes express CXCR2, show reduced mRNA levels of adipogenesis associated genes in CXCR2 KO mice and reduced differentiation in the presence of a CXCR2 inhibitor

Previous in vitro studies have reported that adipocytes themselves express CXCR2 and that it plays a role, in combination with its ligands, in their development and size^24,32,33^. To confirm this expression pattern, we differentiated adipocytes from pre-adipocytes which was confirmed by positive oil red-O staining of the differentiated cells (Figure 5a). qRT-PCR analysis of these cells pre, and post, adipocyte differentiation revealed increased expression of CXCR2 in the differentiated cells (Figure 5b). Quantification of oil red-O staining demonstrated a 30% reduction in the presence of a CXCR2 inhibitor during adipocyte differentiation (Figure 5c), compared to differentiated cells alone, associated with a reduction of PPARγ expression (Figure 5d and Supplementary Figure 1). To extend the in vitro analysis and examine this in vivo, mRNA was extracted from subcutaneous adipose tissue as well as from the tricep, perigonadal and inguinal fat depots of wild type and CXCR2 KO female mice. Non-quantitative PCR (Figure 5e) revealed that cells in all the wild type fat depots expressed CXCR2 mRNA and that it was completely absent in the CXCR2 KO mice. Furthermore, qRT-PCR analysis of these same samples demonstrated no significant differences in the relative level of CXCR2 mRNA expression between these different adipose depots (Figure 5f).

**Figure 5.**
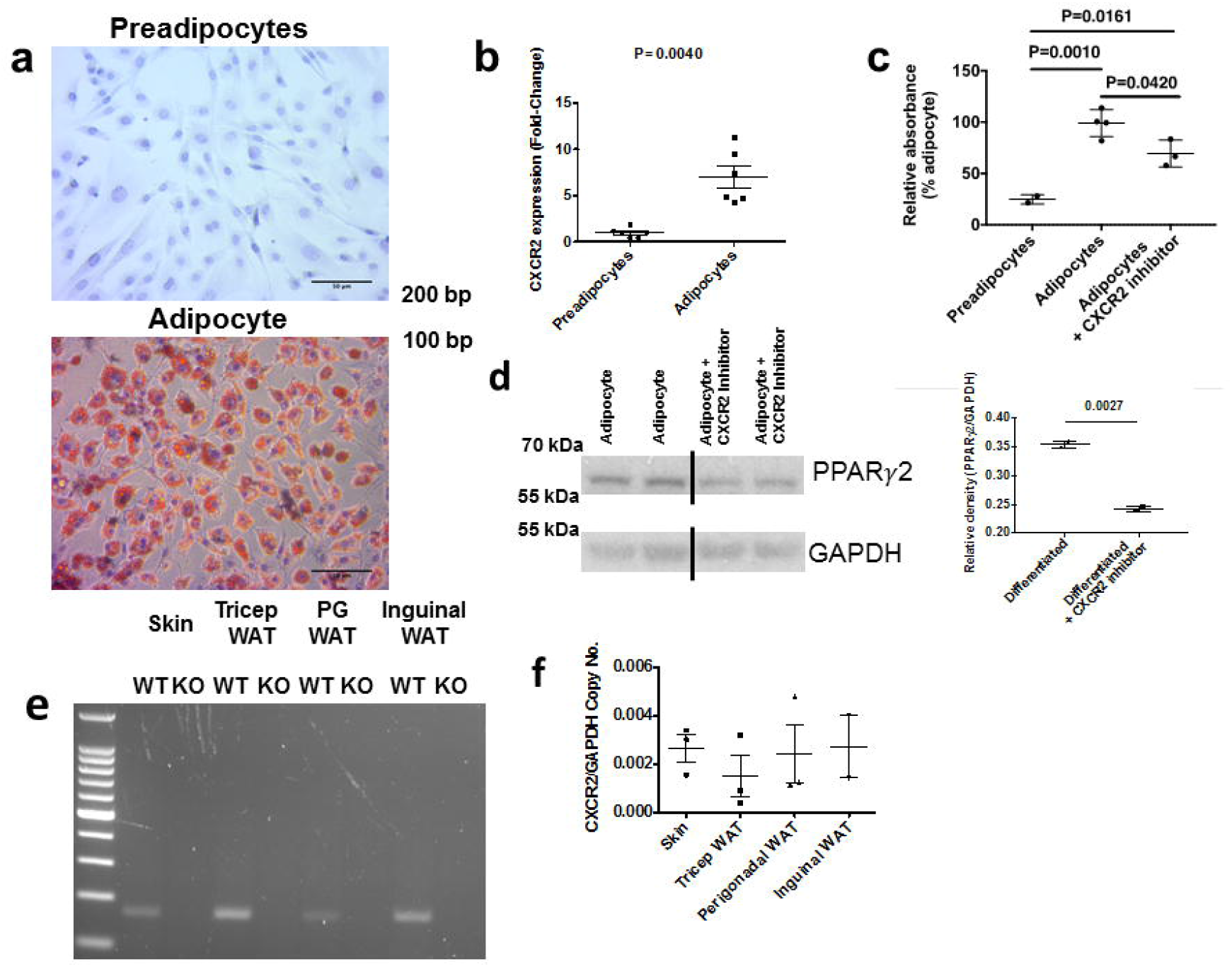
Differentiated adipocytes and fat depots express CXCR2. (a) 3T3-L1 cells were differentiated into adipocytes and cells stained with oil red-O (scale bars-50μm). (b) mRNA was extracted before and after differentiation, reverse transcribed to cDNA and analysed for the CXCR2 expression relative to the house-keeping gene 18s. (c) Oil red-O staining was quantified in undifferentiated and differentiated adipocytes in the absence or presence of a CXCR2 inhibitor (expressed relative to differentiated cells). (d) Adipocytes differentiated in the absence and presence of a CXCR2 inhibitor were analysed for PPARγ protein expression relative to GAPDH levels (left panel) and quantified relative to GAPDH using densitometry. (e) Skin and different adipose depots were dissected from WT and CXCR2 KO mice and analysed for CXCR2 expression by non-quantitative PCR as demonstrated by an expected product of 169 base pairs from the CXCR2 specific primer set. (f) Quantitative real-time PCR was used to analyse CXCR2 expression in skin or adipose tissues, relative to the house-keeping gene, GAPDH. Data are plotted as mean (± SEM), where each symbol represents an experimental replicate (a, b, c and d) or an individual mouse (f). Analysed using an unpaired T test with Welch’s correction (b and d) or one way ANOVA with TUKEY’s multiple comparison test. Data are representative of two separate experiments.

### Adipogenesis-related genes are downregulated in adipose tissue of CXCR2 KO mice

qRT-PCR was next used to investigate relative mRNA expression of PPARγ and fatty acid binding protein 4 (FABP4) in inguinal and subcutaneous adipose depots. Both of these genes play a key role in adipogenesis and have been shown, in vitro, to be transcriptionally regulated in response to CXCR2 signalling^24^. In adult mice (8-12 weeks of age) no significant differences in expression of these genes was observed in either of these adipose depots (Figure 6a). However, when mRNA was extracted from the same adipose depots of juvenile (6 weeks of age) female mice and analysed (Figure 6b) we found that FABP4 showed a small, but significant (P=0.0246), reduction in expression in the inguinal fat depot of these juvenile mice and that there was a 50% reduction in the levels of PPARγ mRNA in the skin of juvenile mice (Figure 6b). Interestingly analysis of inguinal adipocytes from juvenile mice demonstrated that there was no significant difference in adipocyte size between wild type and CXCR2 KO females (Figurer 6c). In fact, inguinal adipocytes from wild type juvenile mice (6 weeks old) were smaller than those in adults, whilst there was little difference in adipocyte size between 6 and 8 weeks in the CXCR2 KO mice (Figures 2 and 6c). This suggests that at around 6 weeks of age adipocytes in the inguinal fat depot increase in size during transition to adulthood, a time period key to inguinal mammary gland maturation^12^, and it is at this stage that CXCR2 may play a role in inguinal adipogenesis. This would possibly confirm findings from a previous study demonstrating a role for CXCR2 in adipogenesis at specific points of maturation^24^.

**Figure 6.**
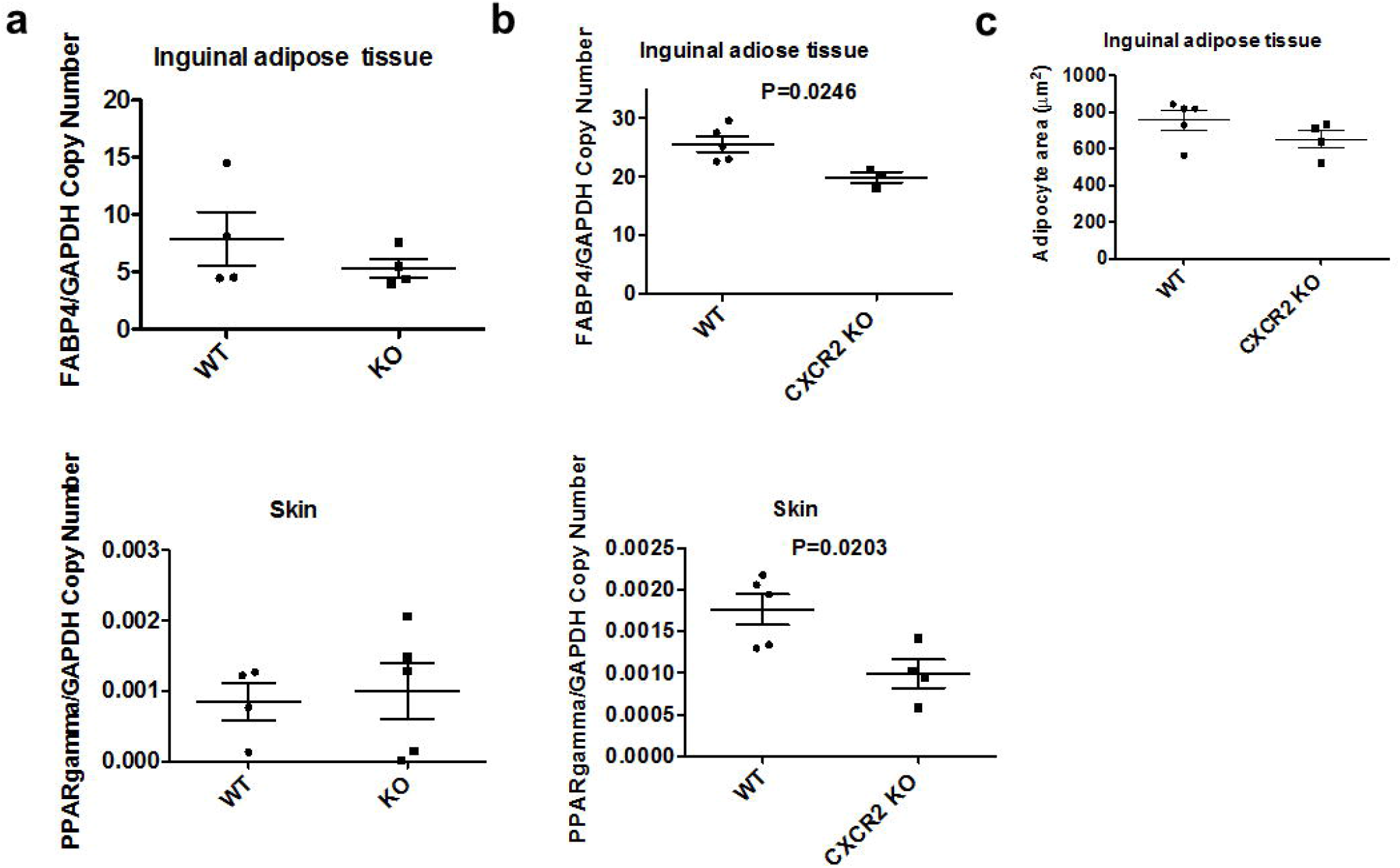
Adipogenesis related genes are down-regulated in the skin and inguinal adipose tissue of juvenile CXCR2 KO mice. mRNA was extracted from inguinal adipose tissue and the skin of (a) adult and (b) juvenile female wild type and CXCR2 KO mice and analysed for expression of PPARγ and FABP4 relative to the house-keeping gene GAPDH. (c) Inguinal adipose tissue from juvenile mice was dissected, fixed, processed and sections stained with haematoxlyin and eosin. The area occupied by individual adipocytes was calculated and plotted. Data are plotted as mean (± SEM) where each symbol represents data from an individual mouse and analysed using an unpaired t test. Data are representative of two separate experiments.

Taken together these data demonstrate that CXCR2 KO mice show a significant reduction in expression of genes that are key to adipogenesis at 6 weeks of age in specific fat depots. The specificity and timing of adipogenesis-related gene expression suggests that CXCR2 is involved in the development of different adipose depots at specific phases of mouse development.

These data, in combination with previous in vitro mechanistic studies^24,29,32^, demonstrate that CXCR2 plays a key role in adipogenesis and regulates adipocyte cell and depot size in vivo, possibly via its direct expression on adipocytes.

## Discussion

In this study we have demonstrated a role for the chemokine receptor CXCR2 in adipocyte cell development and thus adipose depots, in vivo. Our data, in combination with previous detailed mechanistic in vitro^24^ and in vivo^29^ studies, suggest that the role of CXCR2 in resting adipose development is due to its expression on adipocytes themselves, and is therefore independent of neutrophils and their well-characterised pro-inflammatory functions^1^. Whilst the full range of adipogenesis related genes controlled by CXCR2 signalling, and the phase of maturation at which they function, remain to be determined, it is clear that chemokine signaling contributes to the regulation of adipogenesis. This may be in a cell/context specific manner^24^ mediated through ERK and JNK phosphorylation^18^ and by the regulation of expression of adipogenesis-related genes such as PPARγ and FABP4.

This CXCR2 adipogenesis function represents a novel, and non-inflammatory, role for this chemokine receptor and widens our understanding of its role in biology. Several other chemokine receptors have been associated, both directly and indirectly, with regulation of adipose tissue. CCR2 and CX3CR1 have been implicated in adipose regulation via their effects on macrophage infiltration into adipose tissue and the subsequent regulatory role of these cells^30,31^. CXCR4 is expressed on adipocytes and limits obesity, demonstrated by CXCR4 adipocyte-specific knockout mice displaying exaggerated high fat diet-induced obesity, compared to wild type mice^34^. This study suggested that CXCR4 played a role in limiting inflammatory cell infiltration into adipose tissue but also in the thermogenic activity of brown adipose tissue. Additionally, CXCR7 activation has been shown to limit atherosclerosis by regulation of blood cholesterol^35^. Thus CXCR2 represents one facet of chemokine mediated control of adipose tissue and related diseases, however it seems to function at rest via direct adipogenesis effects and not through regulation of inflammatory cells entering into adipose tissues.

This significant effect on a fundamental process such as adipocyte development may provide mechanistic clues to previous phenotypes associated with CXCR2 KO mice. CXCR2 and its ligand CXCL5 are enriched in human atherosclerotic coronary arteries, where a CXCL5 genetic variant may be a molecular marker and target for treatment in coronary artery disease^36^. Furthermore, CXCR2 KO mice have improved sensitivity to insulin in an obesity induced model of insulin resistance^25^. Given the significant role that adipocytes play in atherosclerosis and obesity induced insulin resistance^37^ it seems possible, though speculative at this stage, that CXCR2 mediated adipocyte differentiation may play a significant role in these processes.

In summary we have shown that CXCR2 KO mice have smaller subcutaneous fat depots due to smaller individual adipocytes, a finding similar to adipocytes from other adipose depots. This seems to be due to reduced expression of key adipogenesis-related genes at specific time points in the life cycle of the mouse. Our study, therefore, reveals an atypical function for CXCR2 and contributes to our overall understanding of the regulation of adipocity by chemokines and their receptors.

## Acknowledgements

Work in GJG’s laboratory is funded by a Wellcome Trust Senior Investigator Award and an MRC Programme grant. GJG is recipient of a Wolfson Royal Society merit Award.

## Author Contributions

DPD, JBR, CK, LMR and FS performed experiments. GJG and DPD conceived the study. GJG and DPD wrote the manuscript. All authors approved the manuscript prior to submission.

## Materials and Correspondence

Correspondence and material requests should be submitted to Prof. G Graham or Dr Douglas P Dyer.

